# FUSE: FUsing EEG–MEG in a Shared Embedding via self-supervised learning for BCI

**DOI:** 10.64898/2026.09.09.750152

**Authors:** Giovanni Messuti, Silvia Scarpetta, Pierpaolo Sorrentino, Marie-Constance Corsi

## Abstract

**Objective:** Combining complementary neurophysiological modalities offers a promising strategy for improving motor imagery (MI) brain-computer interfaces (BCIs), but learning shared representations across modalities remains largely unexplored. Here, we propose a two-phase deep learning framework for multimodal EEG–MEG decoding that explicitly decouples representation learning from downstream classification.

**Approach:** In the first phase, a convolutional encoder-decoder learns a shared latent representation by predicting the power spectral density (PSD) of EEG and MEG signals directly from time-domain activity, rather than using the conventional objective of reconstructing the input signal. In the second phase, the encoder is frozen, and its learned representations are reused, without further adaptation, to perform the classification of the downstream MI-BCI task. The framework is evaluated on simultaneous EEG and MEG recordings from 20 participants.

**Main results:** The learned representations consistently outperform conventional handcrafted spectral features, increasing median classification accuracy from 0.734 to 0.794. The multimodal framework also improves performance over MEG alone (median accuracy from 0.680 to 0.794) and yielded a modest increase over EEG alone (median accuracy from 0.765 to 0.794), suggesting potential benefits of multimodal integration over single-modality approaches. Furthermore, the learned latent representations prove to be transferable across participants, with more than half of cross-subject configurations performing within 0.01 accuracy of their subject-specific counterparts.

**Significance:** These findings demonstrate that task-agnostic representation learning can capture physiologically meaningful multimodal neural representations that remain transferable across individuals, offering a promising foundation for more robust and reusable BCI pipelines.

## 1 Introduction

Brain–computer interfaces (BCIs) facilitate communication and control via the modulation of neural activity [1, 2]. Despite their potential, widespread clinical deployment is impeded by significant performance variability, with 15–30% of users exhibiting “BCI inefficiency”, the inability to achieve reliable control despite rigorous training [3]. Current mitigation strategies generally fall into two categories: *machine-centered* approaches, which optimize data quality and classification algorithms [4], and *user-centered* approaches, which adapt to cognitive profiles and neurophysiological markers [5, 6].

A promising yet underutilized avenue lies in enriching input information through multimodal integration. By fusing complementary data sources, *hybrid BCIs* (combining neuroimaging with peripheral biosignals) [7, 8, 9] and *multimodal BCIs* (integrating multiple neuroimaging techniques) [10, 11, 12] aim to robustly decode user intent. Specifically, the combination of Electroencephalography (EEG) and Magnetoencephalography (MEG) offers distinct advantages due to their complementary sensitivities to source depth [13], tissue conductivity [14, 15, 16, 17], and dipole orientation (i.e., radial vs. tangential) [18, 19]. Previous studies have compared BCI performance using EEG and MEG separately [20, 21] or demonstrated that EEG–MEG fusion improves decoding performance [22], highlighting the potential of combining complementary neurophysiological information.

In parallel, Deep Learning (DL, [23]) has emerged as a powerful tool in various BCI applications [24, 25, 26, 27], learning neural representations [28] directly from electrophysiological signals, reducing the dependence on manually engineered features and allowing the extraction of hierarchical patterns from raw data. DL architectures have shown promising results for motor imagery decoding by automatically capturing spatial, temporal, and spectral characteristics of neural activity [29]. More recently, representation learning paradigms, particularly self-supervised approaches, have attracted increasing attention because they aim to learn informative latent representations independently of a specific downstream objective [30]. Despite these advances, the application of representation learning to multimodal EEG–MEG data remains largely unexplored.

In our work, we propose FUSE, a DL-based EEG–MEG fusion framework that learns a shared latent representation in a task-agnostic way, capturing the complementary information carried by both modalities. Unlike many DL approaches, which directly optimize a downstream classification objective in an end-to-end manner, our method explicitly decouples representation learning from task optimization. Rather than learning features tailored to a specific motor imagery task, we first investigate whether a shared latent representation of EEG and MEG signals can be learned, without using information about the labels of the downstream BCI application. Crucially, rather than relying only on the input time series, the representation is learned by predicting their spectral contents, thereby constraining the latent space to preserve physiologically relevant oscillatory information. Only after this representation has been learned, the classification is performed, allowing us to evaluate its intrinsic quality, usability, and transferability as a neural embedding. By separating representation learning from BCI task decoding, the proposed framework aims to learn features that are not tied to a specific problem, potentially enabling their reuse across different BCI paradigms and multimodal neurophysiological applications.

## 2 Materials and Methods

### 2.1 Data and Experimental Protocol

In this study, we used simultaneously acquired EEG and MEG recordings from twenty healthy, right-handed adults (27.5*±*4.0 years; 12 males), all naive to BCI and without any psychological or neurological disorders. Data were recorded at the Paris Brain Institute (ICM) in Paris [31].

Subjects sat in front of a screen at a distance of 0.9*m*. They were asked to perform a 1D, two-target box EEG-based BCI task [32]. They were trained to control a cursor to hit a target. The target was a grey vertical bar displayed on the right side of the screen and could appear in one of two vertical positions: top (up-target) or bottom (down-target). The cursor moved at a constant velocity along the horizontal axis from left to right, while participants controlled its vertical position. To move the cursor upward and reach the up-target, participants performed sustained Motor Imagery of right-hand grasping (MI condition), whereas to hit the down-target, they were asked to remain at rest (Rest condition). Cursor control was based on *α* and *β* band activity derived from sensorimotor areas.

Fig. 1 shows a schematic of the trial timing. The first second corresponds to the Inter-Stimulus Interval (ISI), during which the screen was blank. The target appeared at second 1 and remained visible until second 7. During this interval, participants were asked to perform the BCI task. The cursor appeared at second 3, in the centre-left part of the screen, and persisted until second 6, providing continuous visual feedback until the reaching of the right part of the screen. At the end of the trial, the target turned yellow for 1 second to indicate a successful attempt, or remained gray otherwise.

**Figure 1:**
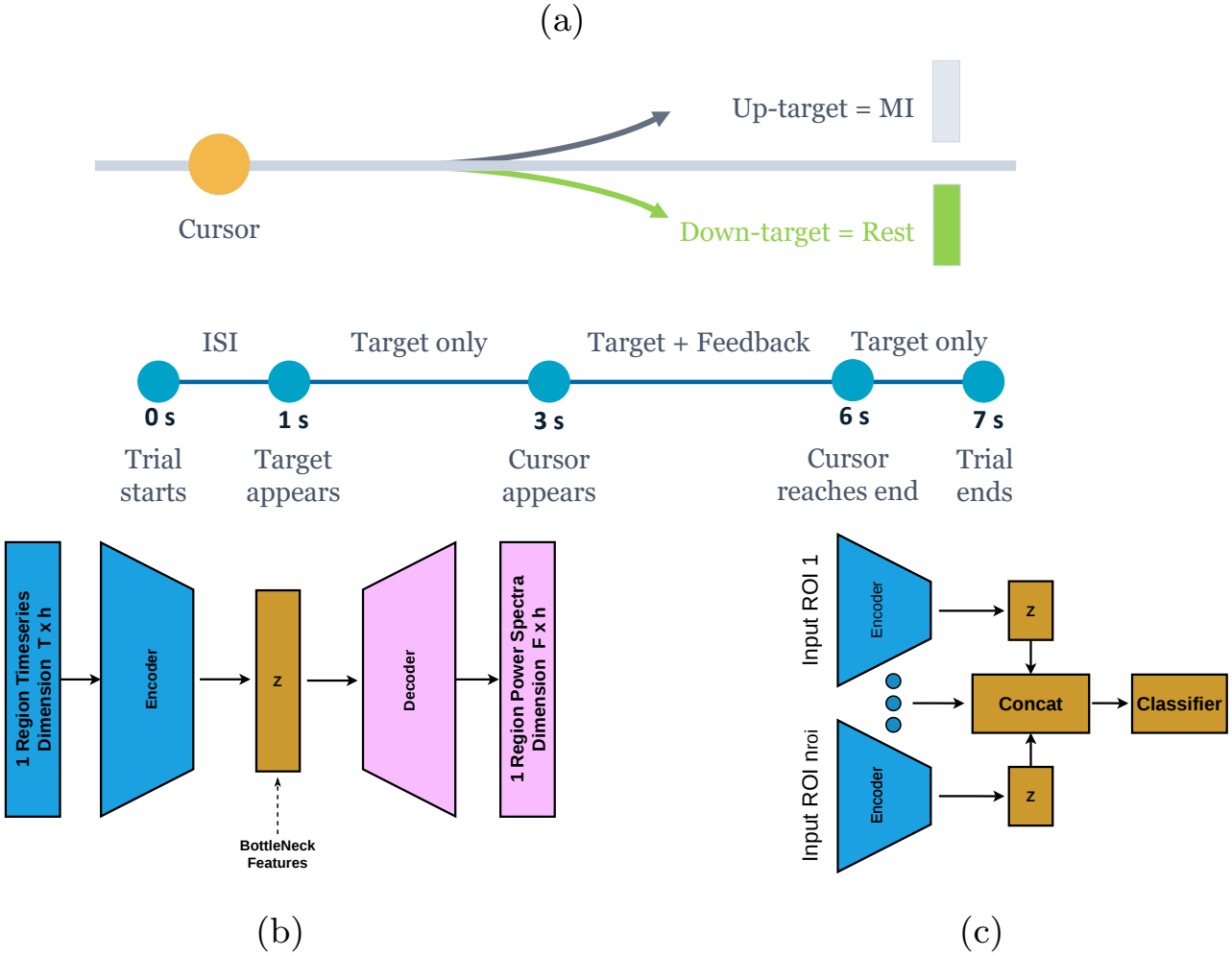
Representation of the methods. (a) - Trial details. Upper part: Scenario encountered by participants in the BCI task. Lower part: Time segmentation of each trial. (b) and (c) present the schematization of the two-phase training framework. (b) - Encoder–decoder network that maps input timeseries (of single ROIs) to their respective power spectrum. The bottleneck latent variables are the features extracted. (c) - Second-phase pipeline. The latent representations of single ROIs are aggregated through concatenation. The same encoder is applied independently to each ROI. A classifier is then placed on top of the feature concatenation. At this stage, the encoder’s weights are fixed.

EEG and MEG signals were simultaneously recorded in a magnetically shielded room with a sampling frequency of 1 kHz and a bandwidth of 0.01–300 Hz. EEG signals were recorded using a 74-channel Easycap system (Ag/AgCl electrodes) arranged according to the international 10–10 system, while the Elekta Neuromag TRIUX MEG system (102 magnetometers and 204 gradiometers) collected MEG data. EEG and MEG signals were source-reconstructed, as described in the original study [31]. Briefly, MEG signals were first denoised using temporal Signal Space Separation (tSSS) [33], and both EEG and MEG data were downsampled to 250 Hz and corrected for physiological artifacts via Independent Component Analysis (ICA) [34]. Source reconstruction was performed using individual anatomical MRI data by means of a Boundary Element Method (BEM) head model [35, 36] and a weighted Minimum Norm Estimate (wMNE) [37]. To perform group analysis, source signals were projected onto a common template (MNI space) [38], and cortical regions were defined according to the Desikan-Killiany (DK) atlas [39], a brain parcellation consisting of 68 Regions Of Interest (ROIs). For further details on experimental design, data acquisition, and source reconstruction, please refer to [31].

For our purposes, we used the last session of the experiment, which consisted of six runs, each comprising an equal number of up-target and down-target trials. We denote by *P* the total number of trials available for a given subject. In general, recorded trials are *P* = 192. However, for some subjects, a small number of trials were excluded due to recording-related issues, resulting in slightly fewer valid trials for these individuals. Across the dataset, the minimum number of valid trials for a subject is 156.

### 2.2 Two-Phase Training Framework

The central idea of this work is to explicitly decouple representation learning [28, 23] from classification. We propose an automatic feature extraction deep learning network, schematized in Figures 1b, and test its ability to provide meaningful features for our BCI problem using the classification pipeline schematized in Figure 1c. The proposed framework consists of two distinct training phases. The first phase, detailed in sect. 2.2.1, is exclusively devoted to representation learning. This phase maps time series of brain activity to their respective spectral representations. By jointly providing EEG and MEG as input signals and reconstructing both of their spectral representations, the network naturally learns a shared latent space that encodes both modalities. The first phase is trained using single ROIs, independently of their anatomical identity, and is completely unaware of the BCI classification task to be performed. For these reasons, we call this phase single-ROI, ROI-agnostic, and task-agnostic.

The second phase, detailed in sect. 2.2.2, evaluates the learned representation through a downstream classification task. During this phase, a ROI selection strategy is performed (sect. 2.2.3), with the aim of choosing which regions to take into account for the classification.

Details about the trainings are provided in sect. 2.2.4. All the models employed in this study were implemented using TensorFlow [40] and scikit-learn [41, 42]

#### 2.2.1 Phase 1 – Feature Learning

The first phase aims to build an automated single-ROI feature extraction pipeline, which represents the core element of our study. We emphasize that our objective is not to optimize BCI classification performance, but rather to propose an alternative feature extraction approach. Accordingly, at this stage we intentionally exclude all components that are related to the downstream classification task; no information about the MI versus Rest labels is available during this phase. The encoder discovers a shared latent representation solely from the statistical structure of the EEG and MEG signals and their corresponding spectral representations, independent of the downstream decoding task. In this sense, the learning procedure is self-supervised: the training targets are automatically derived from the input signals themselves, without using behavioral or task labels. This allows the proposed pipeline to remain independent of any specific BCI paradigm, not tailored to the task considered in this study, and potentially applicable across different experimental settings.

This phase employs a network with an encoder-decoder structure, where both the encoder and decoder have a convolutional architecture. Convolutional Neural Networks (CNNs) are well suited for time series analysis, as they automatically extract local temporal features through learnable filters [43]. In the context of BCI, CNNs have become a remarkably popular feature-extraction approach, because they can learn temporal and spatial filters directly from raw signals in a data-driven manner [44, 45]. A schematic representation of this stage is provided in Figure 1b.

The encoder *E* takes the inputs **x** in the time domain and outputs values **z** into a compact latent representation:

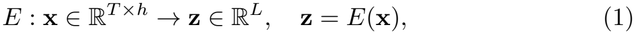

where *L* is the dimension of the latent representation, *T* is the length of the time-series, and *h* the number of different modalities (*h* = 2 if we work with both EEG and MEG; *h* = 1 if we work with a single modality). The input **x** represents the source-reconstructed neural activity, derived from either EEG or MEG recordings, associated with a single ROI, allowing the same feature-extraction pipeline to be applied independently to every brain region. The encoder comprises four convolutional layers with leaky ReLU activation, followed by dropout and pooling layers, and the last layer, representing the bottleneck, is a fully connected layer. For a detailed description of the encoder-decoder architecture, refer to Figure S1.

The decoder *D* maps this representation from the latent space to a target spectral domain output:

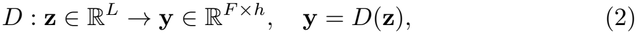

where *F* is the number of frequencies considered. Importantly, the decoder preserves the input structure: the *h* modalities in the time-domain inputs are reconstructed as *h* corresponding spectral profiles. In this way, the network learns to produce a frequency-domain representation for each modality, ensuring consistency between the input and output spaces, while allowing the encoder–decoder network to learn spectral features in a self-supervised manner.

The Decoder comprises a fully connected layer followed by 3 convolutional layers, making use of leaky ReLU activations. The last layer uses a linear activation, as is usual in a regression problem. For the details about the architecture of the encoder and the decoder, refer to Figure S1.

To train the encoder-decoder network, we used the Mean Squared Error (MSE) loss, computed between the PSD estimates of the inputs and the network predictions. We optimized the loss using the Adam algorithm [46] with a learning rate of 0.001, an *ϵ* parameter of 10^*−*4^, and a batch size of 512. We set the maximum number of epochs to 100 and, to prevent overfitting, implemented early stopping [47] that interrupts training if the validation loss does not improve for 10 consecutive epochs.

Unlike standard autoencoders, this latent space is not constrained to preserve all input information; instead, it is optimized to retain only the aspects of the signal that predict its spectral-domain representation. In this way, the network effectively learns an implicit transformation from time-domain neural activity to its spectral representation. The latent space acts as an intermediate computational stage bridging the temporal and spectral domains and encodes all relevant information needed to reconstruct the spectral content.

#### 2.2.2 Phase 2 – Classification

The second phase evaluates, rather than further optimizes, the representation learned during Phase 1. To this end, we perform the BCI classification task using the latent representations learned in the first phase. At this stage, the encoder weights learned in the first phase are kept frozen, without any further adaptation of the representation. This decoupled training strategy allows us to assess the quality of the learned shared latent representation independently of any further optimization of the feature extractor. To further emphasize that the observed performance reflects the quality of the learned latent representation rather than the capacity of the downstream classifier, we deliberately employ a lightweight classifier rather than a deep architecture, thereby keeping the overall complexity confined to Phase 1. This follows the standard practice of evaluating self-supervised representations through a frozen encoder paired with a lightweight linear downstream classifier [48, 49].

Although the feature-extraction network is trained on single ROIs, motor imagery does not arise from isolated brain areas, but from the coordinated activity of distributed cortical regions [50, 51, 52]. To account for this distributed organization and benefit from integrating information across multiple regions, the encoder is applied independently to each ROI to produce one latent representation per region, and the resulting ROI-specific vectors are concatenated into a single feature vector, as shown in Figure 1c. This preserves the information extracted from each region individually while allowing the classifier to leverage complementary information across the brain.

Specifically, we chose to train a Support Vector Machine (SVM) [53] on concatenated latent vectors to discriminate between MI and Rest conditions. This is consistent with its established use as a robust classifier for EEG-based BCI [4, 54], and is particularly useful in settings with limited sample sizes and high-dimensional feature spaces [55].

#### 2.2.3 ROI Selection

The Desikan-Killiany atlas parcellates the brain into 68 regions. The feature-extraction pipeline described in the first phase operates on a single ROI, independent of its anatomical identity. However, to reduce the dimensionality of the feature space and improve computational efficiency, only a subset of the available ROIs is provided to the classification stage depicted in Figure 1c.

Hand-grasping motor imagery predominantly recruits sensorimotor areas, including primary motor and somatosensory areas as well as premotor regions [56, 57, 58], which, under the DK parcellation, correspond to eight cortical ROIs (i.e., precentral, paracentral, postcentral, caudal-middle-frontal, for each hemisphere). Rather than restricting the analysis to this *a priori* anatomical selection, we also account for inter-subject variability in the spatial distribution of brain activity during motor imagery tasks.

To this end, we propose a data-driven ROI selection strategy based on the well-known phenomenon of Event-Related Desynchronization (ERD) [59, 60], which characterizes the modulation of sensorimotor rhythms during motor imagery. Specifically, ROI selection is performed independently for each subject using the spectral content of the signals. For each ROI, each trial, and each of the *h* modalities (refer to eq. 1 for *h*), the power spectral density (PSD) is estimated using the Welch method, which provides a robust estimate of the PSD by averaging periodograms computed over partially overlapping signal segments [61]. The PSD is computed using the `scipy.signal.welch` implementation [62] with parameters `nperseg` = 250, `noverlap` = 125, and `nfft` = 300, while all remaining parameters are kept at their default values. Since the signals are sampled at a frequency sf = 250 Hz, this configuration yields a frequency resolution of approximately sf*/*`nfft` = 0.83 Hz. To capture the time interval in which neuromarkers of motor imagery activity are expected to be most pronounced, PSD estimates are computed exclusively over the last 3 seconds of each trial, corresponding to the period during which continuous visual feedback is provided. Furthermore, the analysis is restricted to the [8, 30] Hz frequency range, as the ERD phenomenon associated with motor imagery is predominantly observed within this interval [63]. Consequently, the PSD of each trial is represented by n_f_ = 27 frequency bins spanning the considered frequency interval. Formally, for a trial represented by the multivariate time series **x**, the PSD associated with modality *i* is computed as:

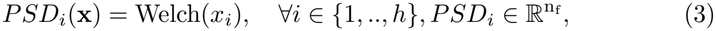

where Welch(·) is the Welch estimate over the last 3 seconds of its argument with the parameters defined above.

Once the PSD estimates are computed, ROI selection is performed independently for each subject. For each ROI, the PSDs corresponding to the MI trials and the Rest trials are separated into two groups. For each modality *i*, and each frequency bin *j*, the difference between the two conditions is quantified using Cohen’s d effect size [64]. Denoting by *r* the ROI index, we have:

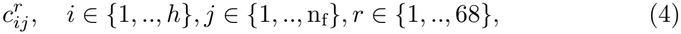

To summarize the discriminative power with a single score, we assigned to each ROI and modality the maximum effect size across the considered frequency range:

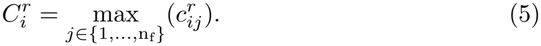

Consequently, each ROI is associated with one score for each modality.

We denote by *n*_roi_ the parameter that determines the number of ROIs retained for the classification stage. In the single-modality setting (*h* = 1), all ROIs *r* are ranked according to their unique score 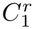, and the *n*_roi_ regions with the highest rank are selected. In the multimodal setting, ROI ranking is performed independently for each modality. The top *n*_roi_ ROIs are selected for each modality, and the final set of ROIs is obtained as the union of the modality-specific selections. We refer hereinafter to the multimodal selection as the Cohen (EEG ∪ MEG) selection.

#### 2.2.4 Training details and signal preprocessing

A separate pipeline was trained for each subject. Model evaluation was performed using five-fold stratified cross-validation [65], ensuring that each fold preserved the original proportion of MI and Rest trials. For each fold, 80% of the trials were used for training, while the remaining 20% constituted the validation set. We wish to highlight that the ROI selection procedure described in Section 2.2.3 is performed exclusively on the training set and imposed onto the validation set, thereby preventing data leakage.

Since the proposed framework consists of two distinct training phases, the validation set was further split into two equally sized subsets, denoted as *Val1* and *Val2*. While the training set was shared by both training phases, the *Val1* subset was exclusively used during the first phase to monitor the training of the encoder-decoder network. The second subset, *Val2*, was reserved for the subsequent classification phase. This separation prevents information leakage between the two stages. In particular, the latent representations used to train and validate the classifier are generated by the encoder whose parameters have been selected without ever observing the samples contained in *Val2*. Consequently, the evaluation of the classification stage remains independent of the model-selection procedure adopted during feature extraction.

Each trial comprises 7-seconds-long signals from 68 ROIs and *h* modalities. To capture the period in which task-related neural activity is expected to be most pronounced, we restrict the analysis to the 3 seconds where the visual feedback is present. The objective of the proposed ROI selection criterion is to identify the most informative brain regions for the downstream BCI classification task, whereas the encoder-decoder network is designed to learn a task-agnostic feature representation. For this reason, no ROI selection is performed before the first training phase, and signals from all 68 ROIs are provided at this stage. As described in Eq. 1, the encoder processes one ROI at a time, irrespective of its identity. Consequently, the first phase learns a generic mapping from the time-domain activity of a single ROI to its corresponding spectral representation, making the feature-extraction pipeline both single-ROI and ROI-agnostic.

Practical BCI systems require low-latency processing in order to provide fast feedback to the user [66]. Accordingly, rather than processing the entire feedback interval as a single input, we test the proposed pipeline on shorter temporal windows. Specifically, the input length in Eq. 1 is set to *T* = 256 samples (approximately 1 s). Before windowing, each trial is band-pass filtered in the [4, 45] Hz frequency range. The 3-s feedback interval is then partitioned into overlapping windows of 256 samples with an overlap of 171 samples. Since seven windows would require a total of 766 samples, and the feedback interval consists of 750 samples, the final window extends 16 samples beyond the feedback interval, incorporating samples from the subsequent period, during which no visual feedback is provided. This pipeline yields 7 windows per trial. The overlap between consecutive windows increases the number of training samples while preserving temporal continuity, effectively acting as a data augmentation strategy.

The number of training samples available during the first phase is:

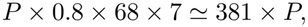

where P is the total number of trials, 0.8 denotes the fraction of trials assigned to the training set, 68 is the number of ROIs, and 7 is the number of windows extracted from each trial. We recall that P varies across subjects, ranging from 156 to 192, with most subjects having the full set of 192 valid trials. This results in a number of training samples ranging from 5.9 *×* 10^4^ to 7.3 *×* 10^4^. The factor 68 arises from the first phase, that processes each ROI independently, i.e., it treats every ROI-window pair as a distinct training sample. In contrast, the second phase operates on the latent representations obtained by aggregating information across ROIs. Consequently, each training sample corresponds to a single trial window rather than to an individual ROI, and the number of training samples is therefore:

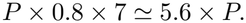

To obtain the spectral targets **y** used to supervise the decoder in Eq. 2, we used the MNE-Python package [67] and estimated the PSD using the multitaper method implemented in `mne.time_frequency.psd_array_multitaper`, with a smoothing bandwidth of 6 Hz and default parameter settings. The method is based on discrete prolate spheroidal sequences (DPSS) tapers [68]. For each input window, PSD estimates are computed over the [8, 30] Hz frequency range, using 24 frequency bins. Because PSD values typically exhibit a 1*/f* -like decay [69], with lower frequencies dominating in magnitude, we take the base-10 logarithm of the PSD estimate to compress the range of variability of the spectrum. The common practice of using the logarithm of the PSD [70, 71] prevents the decoder from being biased toward components at low frequency and encourages it to also capture structure with higher frequencies. The resulting representation corresponds to the **y** used in Eq. 2.

Both the input time windows **x** and the corresponding logarithm of PSD estimates **y** are normalized using the same procedure. An independent per-window normalization would suppress inter-sample amplitude variations that may carry information relevant to the spectral content of the signals. Since these variations may be potentially related to ERD phenomena, preserving them during preprocessing is desirable. Therefore, instead of normalizing each time window (or PSD estimate) independently, normalization is performed jointly across all training samples. Moreover, because different ROIs exhibit distinct amplitude distributions and variability, normalization is performed independently for each brain region. Specifically, for each ROI, the mean and standard deviation are computed over all training samples associated with that region. Each sample is then standardized by subtracting the corresponding mean and dividing by the corresponding standard deviation. The normalization parameters estimated from the training set are subsequently used to normalize the corresponding validation samples.

## 3 Results

A separate two-phase pipeline (described in Section 2.2) was independently trained for each subject. The proposed framework combines three main methodological components: (i) a task-agnostic and region-agnostic encoder-decoder network that learns latent representations from single-ROI neural activity, (ii) the multimodal integration of EEG and MEG signals through a shared latent space, and (iii) a subject-specific ROI selection strategy based on spectral effect sizes.

The experiments presented in this section are designed to assess the contribution of each of these components. Rather than focusing on finding the best classifier to maximize classification accuracy, our objective is to determine whether the latent representation learned during Phase 1 captures meaningful information that can support downstream BCI decodings. For this reason, we used a lightweight classifier when evaluating the downstream decoding. Since the encoder is trained independently of the MI/Rest labels and remains frozen during the classification phase, the resulting classification performance provides an indirect measure of the discriminative information encoded in the learned feature space.

We begin by evaluating the encoder-decoder network trained during the first phase, assessing its ability to reconstruct the spectral representation of the input signals. This analysis provides evidence that the learned latent space captures information relevant to the spectral representation of the input signals. We then examine the contribution of the proposed ROI-selection strategy, multimodal EEG–MEG fusion, and the learned latent features through a comparison with a conventional approach. We end the section with an investigation of how the feature extraction pipeline generalizes in a cross-subject framework.

### 3.1 First phase training evaluation

The first phase aims to train networks capable of learning meaningful latent representations that retain the information necessary to reconstruct the spectral content of the input signals. As described previously, we use input time series of length 256 samples and output representations with 24 frequency bins; i.e., we set *T* = 256 and *F* = 24 in Eqs. 1 and 2, respectively.

A key hyperparameter in designing an encoder–decoder architecture is the dimensionality *L* of the latent space. We determine *L* based exclusively on the ability of the network to reconstruct the target power spectra. This choice ensures that the dimensionality of the latent representation is optimized solely according to the objective of the first phase, without relying on classification performance or any other task-specific information. In this way, the selection of *L* remains consistent with the task-agnostic nature of the proposed feature-extraction stage.

Figure 2 shows the effect of the latent space dimensionality on the MSE loss computed on the validation set, for each modality case. We investigate both the single-modality scenarios (*h* = 1 in Eq.s 1 and 2) and the two-modality scenario (*h* = 2), in which EEG and MEG signals are jointly provided to the network. These settings are denoted as EEG, MEG, and EEG+MEG, respectively. Each scatter point represents a single network, trained on a specific subject for the respective setting. The violin plots represent the distribution of the MSE across subjects, for fixed modality and latent dimension *L*. We chose to investigate *L* ∈ *{*4, 8, 16*}*. Values smaller than 4 would impose a strong compression and result in an overly restrictive bottleneck. Conversely, values larger than 16 were not considered, as they would provide limited compression relative to the *F* = 24-dimensional target representation (in the single modality case). For both EEG and MEG, a latent dimension of *L* = 4 resulted in significantly higher MSE values, while no substantial improvement was observed when increasing the dimension from *L* = 8 to *L* = 16. In contrast, for the combined EEG+MEG modality, using *L* = 16 led to lower MSE values than *L* = 8. This reflects the greater expressivity required in the latent space to model the increased complexity of the multimodal output. Based on these results, we fixed the latent dimension to *L* = 16 for all subsequent analyses, which optimizes the MSE performance for the two-modality case, and we adopted the same value for the single-modality analyses, to maintain consistency across all experiments.

**Figure 2:**
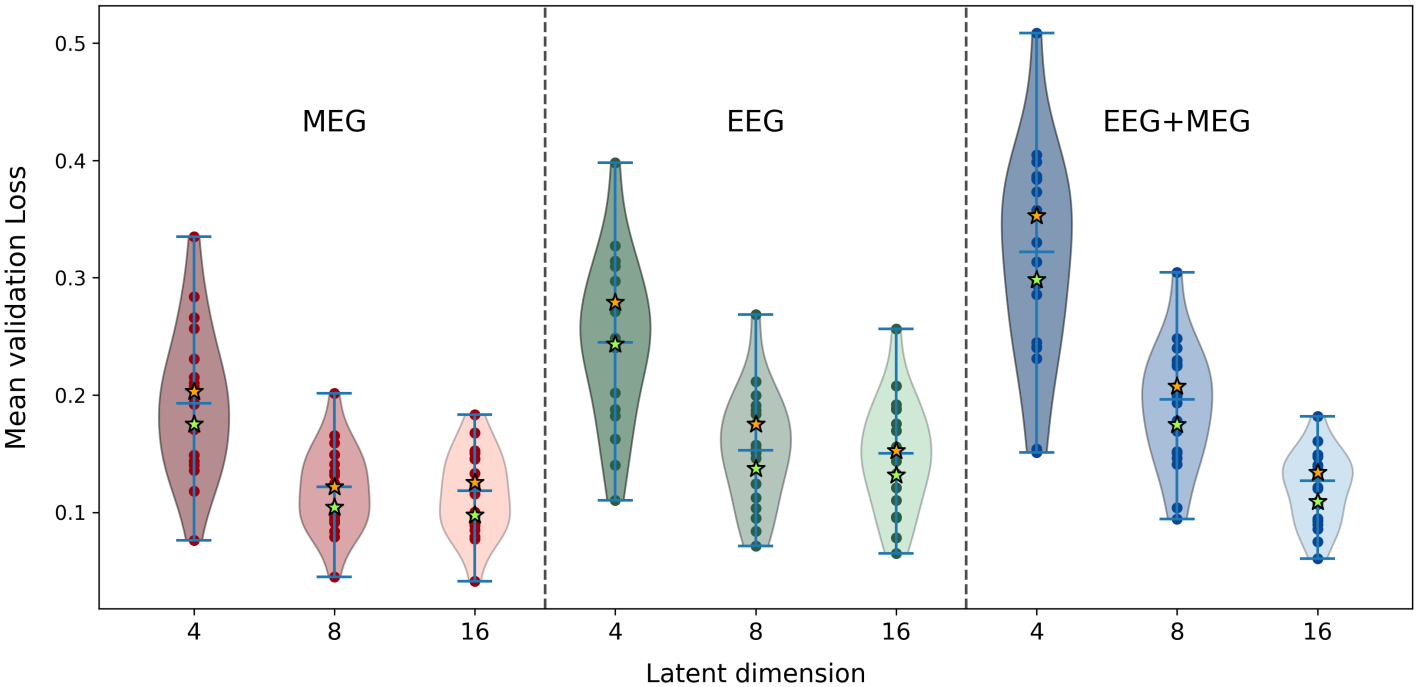
Violin plots showing the distributions of validation loss among subjects varying the latent dimension *L*, for each modality. The orange star identifies subject 2, and the green star identifies subject 13; they are used as references to produce the plot in figure S2, as their validation losses are, on average, the closest to the medians of each distribution, making them suitable representative reference subjects.

Figure S2 illustrates the ability of the different networks, with *L* = 16, to reconstruct the PSD of the input signals, using the multitaper estimates (see Sect. 2.2.4) as targets. Examples are provided for Subjects 2 and 13, represented by orange and green stars, respectively, in Fig. 2. For each subject, the first row shows examples from individual signals, while the second row shows the average reconstruction over the entire *Val1* set. The results highlight that the networks successfully reproduce the spectral profiles across the entire frequency range considered.

### 3.2 Second phase training evaluation

We now focus our attention on the second training phase, in which the BCI classification task is performed. No additional encoder–decoder networks are trained at this stage. Instead, for each subject, we use the networks presented in Sect. 3.1, considering only those with a latent dimension of *L* = 16. The encoder component of each network is extracted, and its weights are frozen. The resulting latent representations are then used as input to SVM classifiers, which are trained during the second phase. In the following sections, we systematically investigate the contribution of each component of the proposed pipeline.

#### 3.2.1 Effect of ROI selection

We first evaluate the proposed ROI selection criterion, providing both neurophysiological insights and an assessment of its impact on the subsequent classification performance. To isolate the effect of the proposed selection strategy, the analysis of this section is conducted in the single-modality setting (*h* = 1 in Eq. 1), using two pipelines for each subject, processing either EEG or MEG signals.

Hand motor imagery is known to predominantly engage the sensorimotor network, which corresponds to eight cortical ROIs in the DK atlas. To enable a direct comparison between this anatomically motivated selection and our proposed data-driven approach (see Section 2.2.3), we set the selection parameter to *n*_roi_ = 8, thereby retaining eight ROIs. We applied the selection procedure separately for each modality (EEG and MEG) and each subject.

Notice that, during the training of the second phase, ROI selection is performed independently within each cross-validation fold to avoid data leakage. Before analyzing the fold-wise selection results, we first inspect the regions identified by the selection procedure when applied to all available trials from each subject, i.e., outside the cross-validation procedure. This will serve as a reference for comparing the fold-wise selections. After selecting the *n*_roi_ ROIs for each subject, we colored each ROI on a brain plot based on the number of subjects for which that region is selected. Figure S3 presents the results of the selection when performed outside the loop, both for EEG and MEG.

Figure 3a summarizes instead the ROI selections obtained within the cross-validation procedure for the EEG modality. Since each subject contributes five selections (one per fold), the contribution of a selected ROI is normalized by the number of folds. Consequently, a region selected in a single fold contributes 0.2 to the displayed value, whereas a region consistently selected across all five folds contributes 1. For reference, the same panel also reports the motor-related ROIs, which are highlighted in yellow. The corresponding results for the MEG modality are reported in Figure S4a.

**Figure 3:**
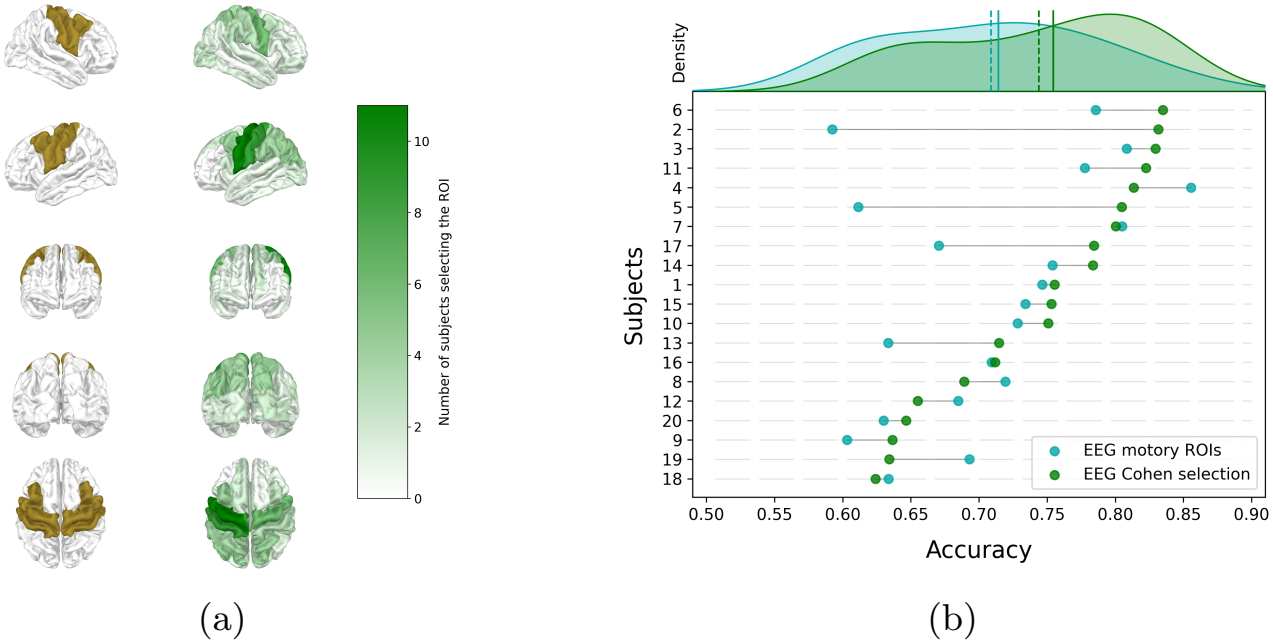
Comparison of ROIs selected based on the Cohen-based selection (*n*_roi_ = 8) with sensorimotor ROIs and performance effects using the EEG modality. Mean accuracy was 0.709 and 0.744, and median accuracy was 0.714 and 0.754, using sensorimotor ROIs and Cohen-selected ROIs, respectively. (a) - Brain plot comparing, in different visualizations, sensorimotor ROIs (in ochre) and ROIs selected by our criterion (in green) within the cross-validation loop for EEG. (b) - Lollipop plot showing the effect of region selection on the performance for each subject. At the top, we show their distributions, along with their means (dashed lines) and medians (solid lines).

Comparing Figure 3a with Figure S3a (and similarly Figure S4a with Figure S3b), we observe that the fold-wise selections and those obtained using all trials are highly consistent. This confirms the robustness of our selection criterion across cross-validation. Furthermore, when comparing the brain plots showing the ROIs associated with the sensorimotor region with the ROIs selected by our method in Figure 3a, the overlap is evident, supporting the neurophysiological validity of our approach. Finally, we note that the left hemisphere is more represented in the selection, which is expected because the MI task in this study involves the right hand. We also observe the selection of regions beyond the sensorimotor areas. However, these additional regions do not show a strong overlap across subjects, indicating that their selection is highly subject-specific. This is visible in the brain plots, where several ROIs appear only lightly colored, meaning they were selected only a few times across subjects.

Figures 3b and S4b compare the classification performances obtained in the second phase when using the eight sensorimotor ROIs or the eight ROIs selected by our criterion, for EEG and MEG, respectively. The central plots summarize subject-wise performance differences between the two frameworks. For some subjects (e.g., 16, 1, 7), performance under the two ROI-selection strategies is comparable in both modalities. In contrast, other subjects (e.g., 2, 13, 17 for both modalities; 5 for EEG; 6 and 8 for MEG) show substantial improvements of 0.1 or more when ROIs are selected according to our criterion based on Cohen’s d effect size. These results clearly indicate that incorporating subject-specific neurosignatures is essential for certain individuals. Above the lollipop plots, we show the distribution of performance, with its mean and median. The improvement obtained when using our ROI-selection criterion is evident. As noted above, the selected ROIs are subject-specific, meaning the method selects different spatial patterns across individuals. This is visible in the variability of selected regions and is consistent with the neurophysiological heterogeneity across subjects. The substantial performance gains (particularly for some subjects) with these individualized ROIs show that a targeted, subject-specific selection strategy is highly beneficial and, for some subjects, necessary to achieve optimal decoding performance.

To assess differences in classification performance, we performed paired two-sided Wilcoxon signed-rank tests comparing performance with sensorimotor ROIs and with Cohen-selected ROIs. For MEG, Cohen-based selection resulted in a statistically significant improvement (*p* = 0.0042), while the corresponding EEG comparison yielded *p* = 0.07. Although the EEG comparison did not reach the conventional significance threshold of *α* = 0.05, the substantial accuracy improvements observed for several subjects (up to 0.1 or more), together with the overall higher performance obtained with the Cohen-based selection, highlight the potential benefit of subject-specific ROI selection.

#### 3.2.2 Effect of two-modalities

In the previous section, we demonstrated how the proposed ROI selection criterion can substantially improve classification performance. We now compare the performances obtained using EEG and MEG independently. For a fair comparison, we used the same cross-validation splits for both modalities, so the only difference between the two pipelines is the input modality. The results, reported in Figure S5a, show that EEG generally achieves higher classification performance than MEG. Nevertheless, MEG can outperform EEG for some subjects (most notably Subjects 7, 13, and 20), suggesting that the two modalities can capture complementary information rather than redundant neural signatures.

Because ROI selection is performed independently for each modality, the selected region sets are not necessarily identical between EEG and MEG. To verify that the observed differences are attributable to the recording modality rather than to different ROI selections, we repeated the comparison using a common ROI set. Specifically, for each subject, we defined a new set of selected ROIs as the union of the EEG- and MEG-selected regions (we introduced this selection in sect. 2.2.3 as the Cohen (EEG ∪ MEG) selection) and trained separate single-modality classifiers using this common set. The corresponding results are shown in Figure S5b and are consistent with those obtained using single-modality ROI selections.

These observations suggest that, although EEG provides superior overall performance, MEG contains, for some subjects, complementary information that can improve the performance. Motivated by this, we investigate the multi-modal framework, where *h* = 2 in Eqs. 1 and 2 and EEG and MEG recordings are jointly provided to the encoder-decoder network. The second-stage SVM classifier operates on the latent representation, with ROIs selected according to the Cohen (EEG ∪ MEG) criterion. We highlight that, in the two-modality setting, the encoder performs, by construction, a multimodal feature fusion by jointly encoding EEG and MEG signals into a shared latent representation. We recall that in this analysis, the latent dimension *L* we used is still equal to 16, as for the single-modality cases.

Figures 4a and 4b compare the classification performance of the proposed multimodal framework with the corresponding single-modality pipelines. To ensure a fair comparison, all configurations use the same ROI selection strategy: Cohen (EEG ∪ MEG) selection. The multimodal framework consistently outperforms the MEG-only pipeline, increasing the mean classification accuracy from 0.688 to 0.780 and the median accuracy from 0.680 to 0.794. We also observe a more moderate but consistent improvement with the EEG-only pipeline, with mean accuracy increasing from 0.768 to 0.780 and median accuracy from 0.765 to 0.794. Moreover, we observe overlap between subjects for which the EEG+MEG pipeline outperforms EEG alone and those for which MEG performs better than EEG. This indicates the importance of the complementary information carried by MEG data relative to EEG alone. The two-sided paired Wilcoxon signed-rank tests comparing the EEG+MEG and EEG pipelines, and the EEG+MEG and MEG pipelines, yielded *p* = 0.14 and *p <* 0.001, respectively. The different statistical outcomes can be interpreted in light of the intrinsic performance difference between the two single-modality pipelines. Because EEG already achieves higher performance than MEG for most subjects, MEG’s additional contribution is expected to be limited. The largest improvements are observed for subjects in which MEG outperforms EEG in the single-modality setting, i.e., those for whom MEG provides particularly informative complementary signals. Since these subjects represent only a subset of the population, the overall improvement of EEG+MEG over EEG, although positive, does not reach statistical significance. This result should not be interpreted as evidence against multimodal fusion. Rather, it suggests that the benefit of incorporating MEG is concentrated in subjects for whom MEG provides particularly informative complementary signals. The multimodal framework therefore offers a more robust approach across subjects, combining EEG’s generally stronger performance with the additional information MEG provides when it is most relevant.

**Figure 4:**
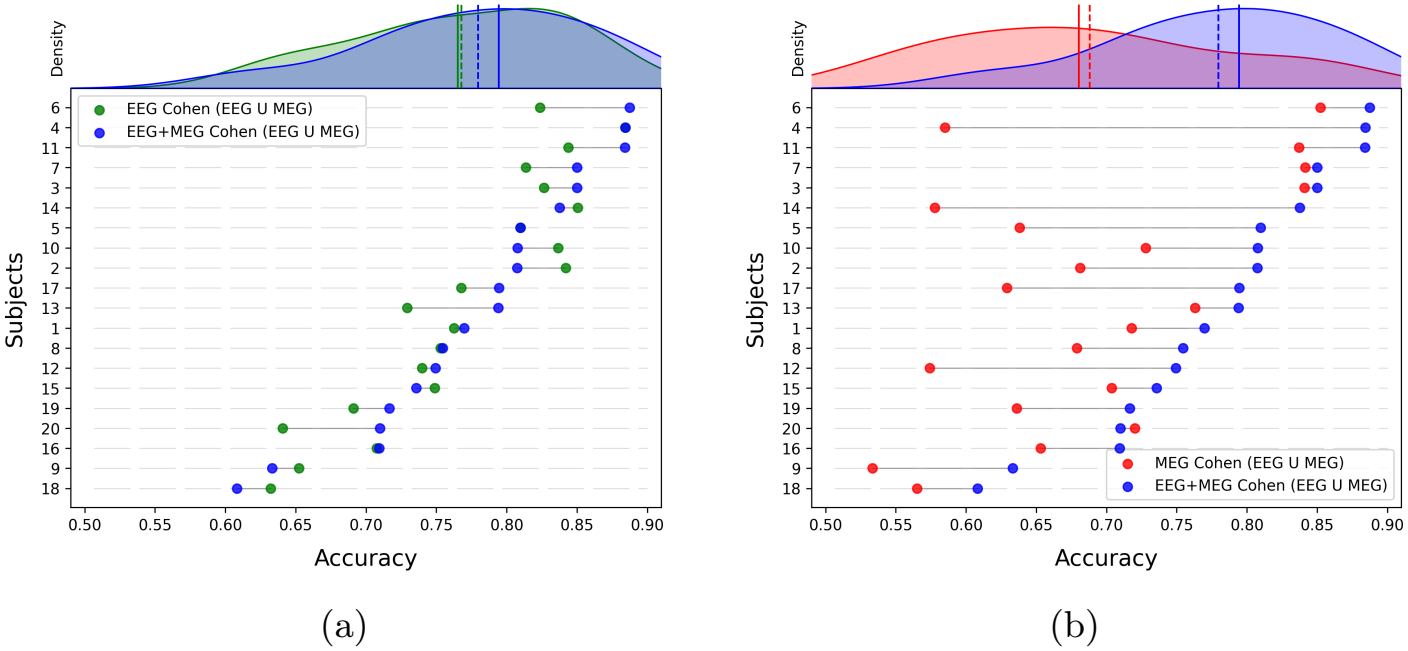
Lollipop plot comparing performances of the two-modality pipeline with the single-modality ones. ROI selection is based on the Cohen (EEG ∪ MEG) selection criterion. Subjects are ordered by performance using the two- modality pipeline. At the top, we show their distributions, along with their means (dashed lines) and medians (solid lines). (a) - EEG modality. (b) - MEG modality.

#### 3.2.3 Comparison with hand-extracted features

Historically, researchers have largely used features derived from PSD estimates in BCI [4]. The only core component of the proposed framework not yet evaluated is the feature-extraction strategy itself. We therefore assess and compare the complete multimodal pipeline to an otherwise identical pipeline in which the learned latent representation is replaced with conventional hand-crafted spectral features.

To ensure a fair comparison, both pipelines use the same cross-validation splits, the same ROI selection strategy (i.e., Cohen (EEG ∪ MEG) selection), and the same SVM classifier. Furthermore, the PSD estimates used to derive the baseline features are the same ones used to supervise the encoder during the first training phase (see Section 2.2.4). Specifically, for each selected ROI and each modality, four features are extracted from the PSD estimate: the mean and the maximum power within the *α* (8–12 Hz) and *β* (13–30 Hz) frequency bands. We then concatenate these hand-crafted features and use them to train the same classifier as in our second phase (see section 2.2.2), replacing the encoder’s latent representations.

Figure 5 shows the improved accuracy of our pipeline compared to a standard feature extraction method. The mean accuracy improves from 0.738 to 0.780, and the median from 0.734 to 0.794. A paired two-sided Wilcoxon test confirms, with *p <* 0.001, the superiority in accuracy obtained using features extracted by our encoder network compared with PSD-extracted features.

**Figure 5:**
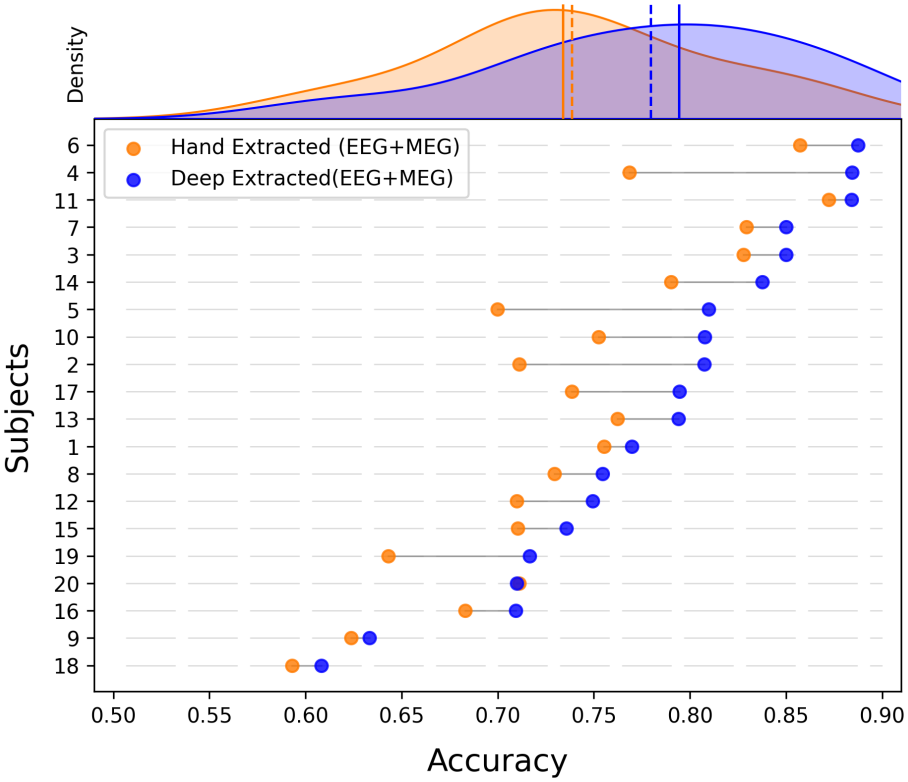
Lollipop plot comparing the effect on performance of the two feature-extraction modalities: encoder-extracted features and classic PSD-based features. ROI selection is based on the Cohen (EEG ∪ MEG) selection criterion. Subjects are ordered based on the performance obtained using our Deep Learning feature-extraction pipeline. At the top, we show their distributions, along with their means (dashed lines) and medians (solid lines).

To isolate the contribution of the proposed feature-extraction strategy from the other components of the pipeline (ROI selection; fusion of two modalities), in Figure S6 we report the comparison between the performances obtained using the representation learned by the encoder and those obtained using conventional PSD-based features in the basic settings, i.e., using a single modality and restricting the analysis to the predefined sensorimotor ROIs. As shown in Figures S6a and S6b, the learned latent representation leads to higher classification performance than conventional PSD-based features for both EEG and MEG.

### 3.3 Cross-subject representation transfer

All previously shown experiments have been conducted in a subject-specific setting, where both the encoder–decoder (first phase) and the classifier (second phase) are trained on data from the same subject. We now investigate whether the learned latent representations are transferable across subjects. Specifically, we evaluate whether an encoder trained on one subject can be used as a generic feature extractor for a classifier trained on a different subject. We conduct this investigation using the two-modality setting with Cohen (EEG ∪ MEG) region selection, whose performance is shown in Figure 5 in blue.

To this end, for each subject *i*, we consider the already trained encoder– decoder network, as described in the first phase. The resulting encoder is frozen and used to extract latent representations from every other subject *j*. For each pair (*i, j*), a second-phase classifier is trained using latent representations extracted by the encoder trained on subject *i*, while the classifier itself is trained exclusively on data from subject *j*. We repeat this procedure for all subject pairs. We will refer to subject *i* as *donor*, as it provides the latent representation, and to subject *j* as the *recipient*, as it receives the space representation and provides the data used to train the classifier. The case *i* = *j* corresponds to the subject-specific pipeline considered in the previous sections.

The results of the cross-subject pipelines are summarized in Figure 6, in a 20*×*20 heatmap. Each row corresponds to a *donor* subject, whereas each column corresponds to a *recipient* subject. To facilitate the comparison, the values in each column are expressed relative to the corresponding subject-specific performance (*i* = *j*). Specifically, each entry represents the difference in classification accuracy between the relative cross-subject setting and the respective subject-specific one in the same column. Rows and columns are ordered according to the average *donor* performance, i.e., the average performance achieved by each encoder across all recipient subjects, allowing us to identify the subjects that produce the most transferable latent representations. The first lines of numbers outside the 20*×*20 grid report the column averages and row averages, respectively. The column averages indicate how much each *recipient* subject benefits, on average, from representations learned from other subjects, whereas the row averages quantify the transferability of each *donor* ‘s learned representation across *recipient* subjects.

**Figure 6:**
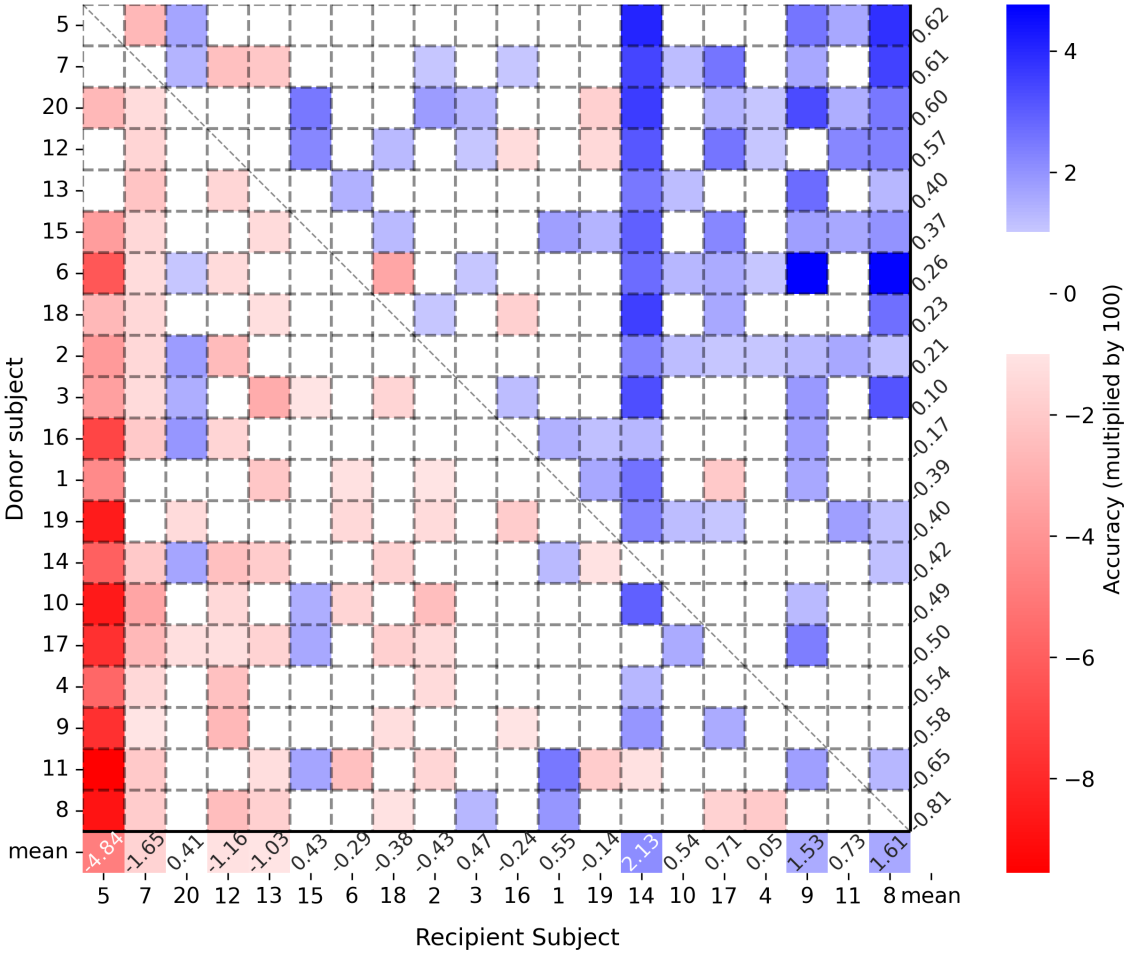
Cross-subject generalization matrix. Each row corresponds to a donor subject, while each column corresponds to a target subject. Each entry reports the difference in classification accuracy between the cross-subject setting and the corresponding subject-specific setting for the target subject (placed on the diagonal of the same column). Values outside the grid report the mean across rows and columns, respectively, quantifying the average transferability of each donor representation and the average benefit each target subject gains from representations learned from other subjects. Values in [*−*1, 1] are shown in white.

The Figure shows the high transferability of the latent representation, as the maximum difference between the cross-subject and subject-specific pipelines never exceeds 0.1. In more than half of the cases (197 out of 380) the difference is smaller than 0.01. We further assessed the relationship between *donor* transfer-ability and *recipient* benefit using Spearman’s rank correlation. The correlation between the row and column averages was -0.81 with a p-value of 1.7 *×* 10^*−*5^.

This strong negative correlation indicates that subjects whose encoders provide the most transferable representations (best *donors*) are the worst *recipients*, i.e., they benefit least from representations learned from other subjects. Conversely, subjects that benefit most from representations learned from other individuals tend to provide less transferable representations themselves. Subject 5 clearly illustrates this pattern: it achieves the highest average donor performance while showing the lowest average recipient performance.

## 4 Discussion

In this study, we introduce FUSE, a two phase, self-supervised deep learning framework for multimodal EEG/MEG decoding that decouples task-agnostic representation learning from downstream classification. Three complementary components contribute to the pipeline: (i) a convolutional encoder–decoder trained to predict the spectral content of neural activity of single ROIs, independent of ROI identity and decoding task; (ii) a shared latent space that integrates EEG and MEG signals into a unified multimodal representation; and (iii) a subject-specific, data driven ROI selection strategy based on spectral effect sizes. The task-agnostic nature of representation learning encourages the encoder to capture intrinsic properties of neural dynamics that, potentially, are reusable across different downstream tasks. In our work, the learned representation is subsequently used, through a lightweight classifier, for MI decoding.

A central aspect of the proposed framework is its representation learning objective. Most existing representation learning approaches rely on autoencoders performing signal reconstruction to learn latent representations [30]. In contrast, our framework learns latent representations by predicting the PSD of the input rather than reconstructing the temporal signal itself. Consequently, the encoder is not required to preserve all the information contained in the input waveform, but instead to retain the information necessary to recover its spectral characteristics. Since specific frequency bands (such as in the *α* and *β* bands) represent one of the most robust neurophysiological signatures of motor imagery [59, 72], this learning objective naturally encourages the latent representation to preserve physiologically relevant information, reducing the sensitivity to temporal fluctuations that are less relevant for motor imagery decoding. In this sense, the proposed framework learns representations that explicitly bridge temporal neural dynamics and their spectral characteristics, rather than simply compressing the input signal.

The comparison with conventional PSD-based features provides further insight into the origin of the performance improvement. Both approaches rely on the same underlying oscillatory information but differ substantially in how this information is represented. Many conventional motor imagery-based BCI pipelines typically summarize oscillatory activity through spectral descriptors extracted from predefined frequency bands [4], whereas the proposed encoder learns a compact nonlinear representation directly from temporal neural dynamics under the constraint of predicting the PSD. The superior decoding accuracy therefore suggests that the learned latent representation is able to capture relationships between temporal and spectral information that are not fully described by handcrafted band-power features while remaining closely linked to physiologically interpretable oscillatory activity.

Rather than assuming fixed, *a priori* spatial patterns for all participants, the proposed approach allows ROI selection to adapt to subject-specific patterns of neural activity. Although the selected regions consistently include the expected sensorimotor cortices, additional cortical areas are identified in a subject-dependent manner, reflecting the inter individual variability in BCI contexts [30]. The various performance comparisons between settings making use of our Cohen-based selection procedure and fixed ROIs selection, highlighted the substantial performance improvements of the proposed selection strategy and demonstrated the value of accounting for subject-specific neural patterns.

The multimodal analyses emphasize the complementary nature of EEG and MEG. While EEG achieved the highest average classification accuracy across participants in a single modality setting, notably, MEG provided better performance for some individual subjects, indicating that the two modalities capture partially distinct aspects of the underlying neural activity. The shared multi-modal latent representation proposed here provides a unified encoding of EEG and MEG signals, allowing complementary temporal and spectral information to be integrated before the classification stage. This integration contributes to the improvement observed for multimodal decoding and is consistent with the complementary biophysical sensitivity of EEG and MEG, which provide distinct but synergistic measurements of cortical electrophysiological activity [13, 14, 15, 16, 17, 18, 19].

The cross-subject analysis provides additional evidence that the proposed representation learning captures properties of neural activity that are not exclusively tied to the subject on which the encoder was trained. Across all donor-recipient pairs, replacing the subject specific encoder with one trained on a different individual resulted in only limited performance difference, with more than half of the cross-subject configurations differing from the subject specific baseline by less than 0.01 in classification accuracy. This finding suggests that the first phase learns a representation that can be transferred across individuals while preserving most of its discriminative utility. From a practical perspective, this property is particularly valuable. Since the encoder can be frozen after representation learning, deploying the framework on a new subject does not require retraining the entire pipeline. Instead, a previously trained encoder can be reused as a generic feature extractor, while only the second phase lightweight classifier is adapted to the new subject. This substantially reduces the computational cost and data requirements associated with calibration burden while remaining consistent with the task-agnostic philosophy of the proposed framework.

Our results should be interpreted in light of some limitations. The cohort is relatively small (20 participants). Moreover, the multimodal pipeline in its current form depends on MEG acquisition, which requires expensive, non-portable, magnetically shielded instrumentation. Emerging optically pumped magnetometer (OPM-MEG) [73, 74] systems are beginning to relax this constraint, but translating the complete EEG–MEG framework into routine use remains an open challenge.

Overall, the present results suggest that PSD prediction represents an effective, task-agnostic objective for learning neural representations from multimodal EEG-MEG signals. Although the present study focuses on a single motor imagery paradigm, the absence of task-specific supervision during representation learning suggests that the same pipeline could potentially be extended to similar BCI paradigms. More broadly, these results indicate that future advances in BCI may depend not only on increasingly sophisticated classifiers, but also on the development of physiologically grounded representation learning objectives.

## 5 Conclusion

This study introduces FUSE, a two-phase self-supervised learning framework for multimodal EEG–MEG decoding that separates task-agnostic representation learning from downstream classification. Unlike other unsupervised feature learning pipelines, it learns a representation that bridges the temporal and spectral domains.

The proposed pipeline addresses several genuine gaps in the current literature: although EEG–MEG fusion has been shown to improve BCI performance, learning a shared representations remains largely uninvestigated [22]; self-supervised approaches to BCI decoding remain relatively underexplored [30]; the few studies that jointly leverage time and frequency domain characteristics of signals typically require frequency information as an explicit input to the model [75]. In our case, an encoder is optimized to predict the spectral content of the input during the first phase, but at inference time, the network does not require spectral information, as it operates directly on time domain signals. This is both a methodological novelty and a practical computational benefit at inference time. The second phase leaverage the features learned in the previous phase, performing classification with a simple lightweight classifier. The learned representation improves motor-imagery decoding in comparison to conventional handcrafted spectral features. In addition, multimodal integration provides more robust performance across participants, improving decoding performance compared to both EEG- and MEG-only pipelines. A subject-specific ROI selection strategy highlighted the importance of accounting for neural variability between individuals and improved decoding performance. Importantly, the latent representation also shows transferability across subjects, suggesting that encoders learned from one individual can potentially be reused as feature extractors for others.

More broadly, our findings support the view that task-agnostic self-supervised representation learning is a promising alternative to standard features for BCI applications [28, 30] for developing more transferable and reusable multimodal BCI pipelines.

## Acknowledgments

This research was supported by the Italian National Recovery and Resilience Plan (PNRR) — European Union — Next Generation EU, DM 118/2023, CUP D42B23001820006 which funded the PhD fellowship of Giovanni Messuti, and by the French National Research Agency (MANET project, ANR-25-CE33-7747-01).

## 6 Supplementary

**Figure S1:**
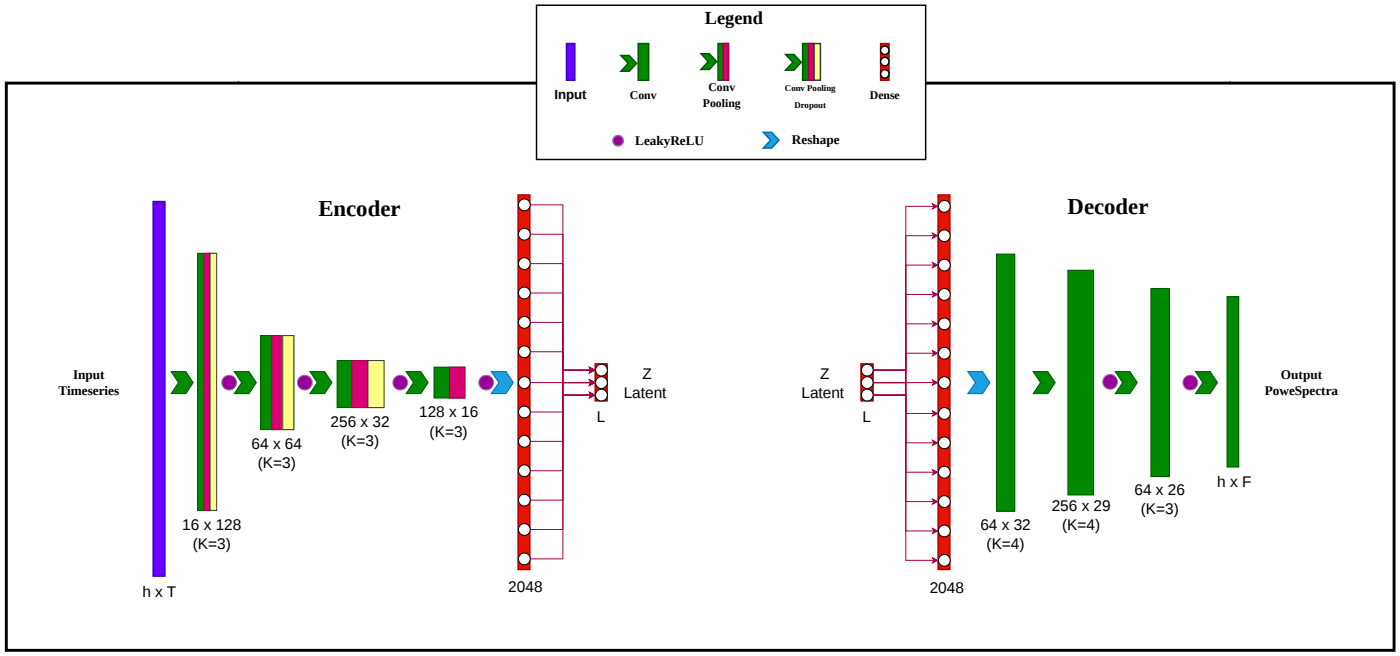
Architecture of the encoder-decoder network. Both the encoder and decoder employ a feedforward convolutional architecture. Each layer is represented as a rectangle, with its type indicated in the legend. The numbers below each layer indicate the corresponding output dimensions, while values in parentheses (*K* = *·*) denote the kernel size of convolutional layers. All pooling operations are implemented using average pooling, and dropout layers use a dropout rate of 0.15. Padding is applied in the encoder convolutional layers to preserve power-of-two dimensions. No padding is used in the decoder. The encoder (left) consists of four convolutional layers followed by a fully connected layer, whose output defines the latent representation, the feature extracted. The decoder (right) receives this latent representation and estimates the spectral representation of the input time series through one fully connected layer followed by three convolutional layers.

**Figure S2:**
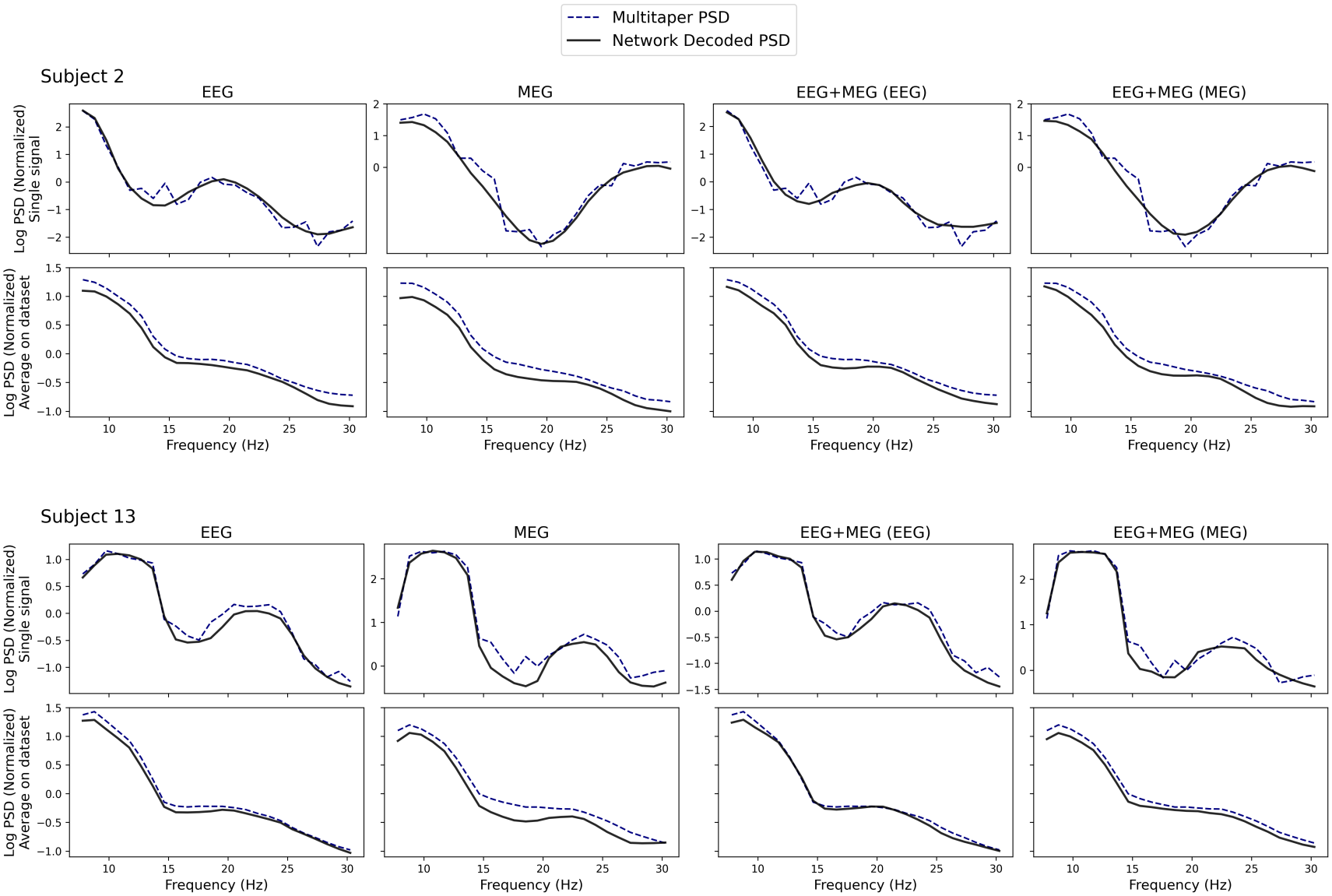
Ability of the encoder–decoder networks to reconstruct the target PSD. For conciseness, only Subjects 2 and 13 are shown, occupying the upper and lower halves of the figure, respectively. The first two columns show the reconstructions obtained from the single-modality networks, while the last two columns show the EEG and MEG reconstructions, respectively, obtained from the (same) two-modality network. For each subject, the first row shows an example of an individual PSD reconstruction, while the second row shows the average PSD across the *Val1* set, comparing the multitaper estimates with the corresponding network reconstructions.

**Figure S3:**
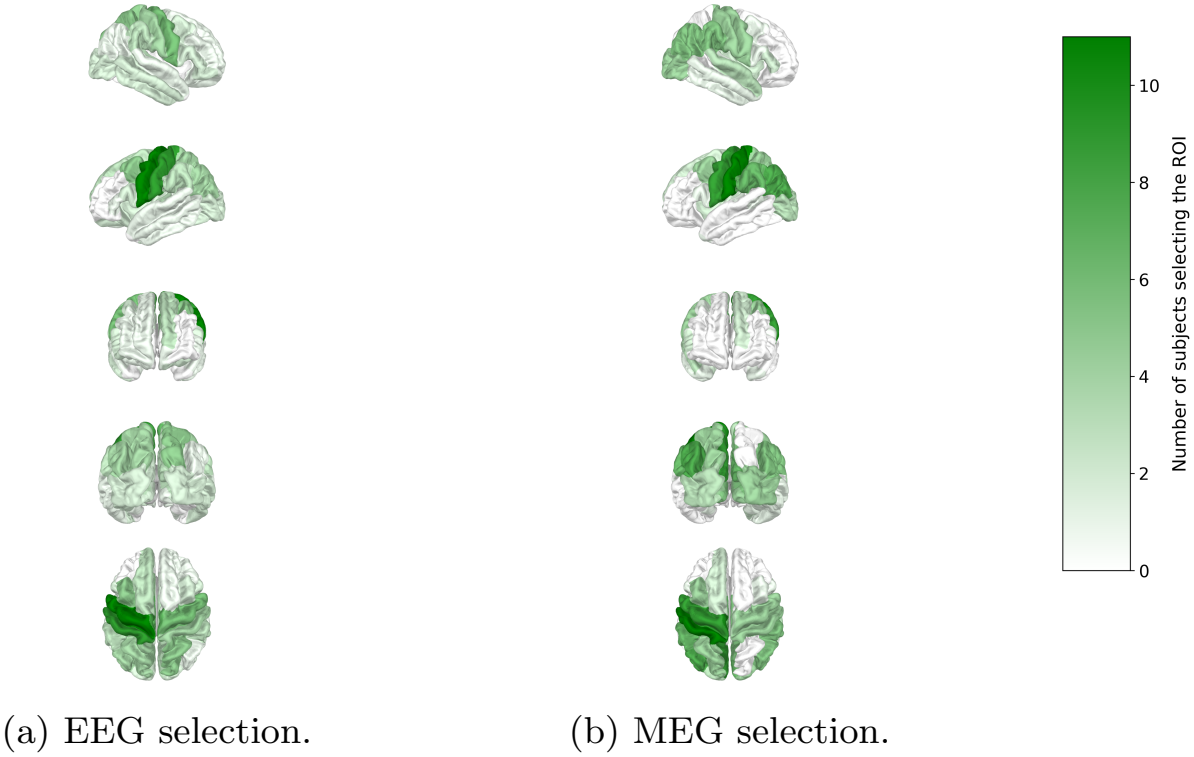
ROIs selected by our criterion based on Cohen’s d effect size, making use of all the data for each subject. The selection is performed independently for each subject. Each brain ROI is colored based on the number of subjects for whom that ROI is selected. The selection is performed both for EEG and MEG modalities, independently.

**Figure S4:**
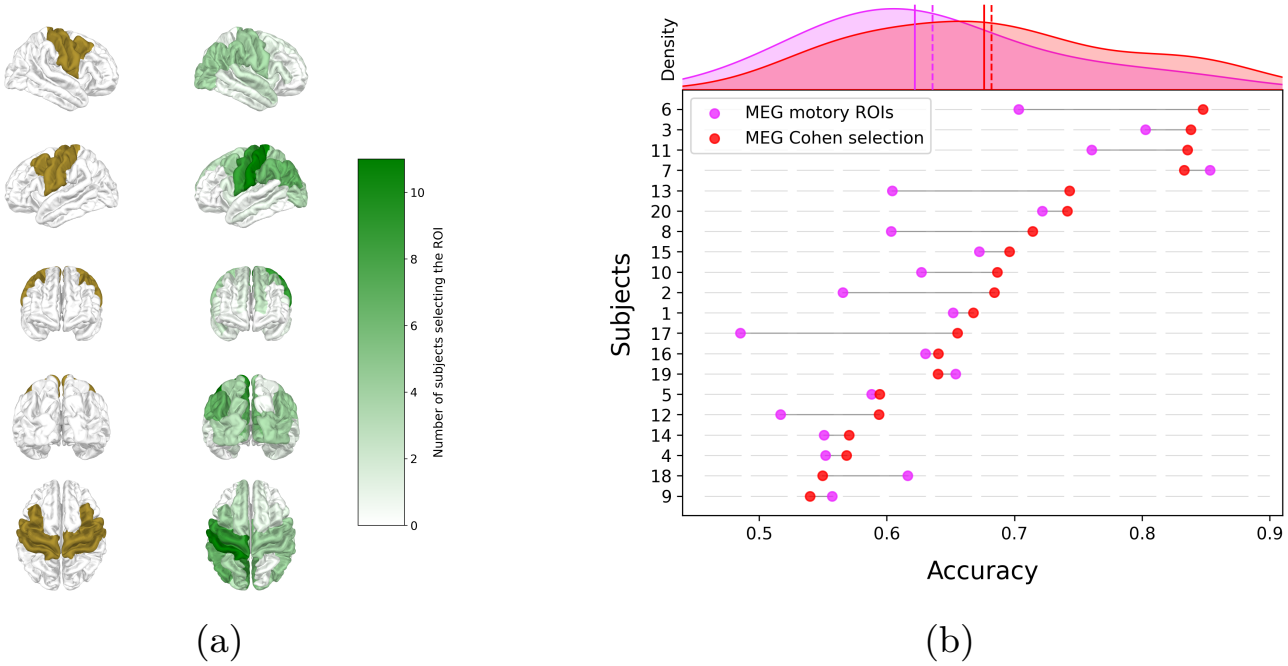
Comparison of ROIs selected based on the Cohen-based selection (*n*_roi_ = 8) with sensorimotor ROIs and performance effects in the case of the MEG modality. Mean accuracy is 0.636 and 0.682, and median accuracy is 0.622 and 0.676, using sensorimotor ROIs and Cohen-selected ROIs, respectively. (a) - Brain plot comparing, in different visualizations, sensorimotor ROIs (in ochre) and ROIs selected by our criterion (in green) within the cross-validation loop for MEG. (b) - Lollipop plot showing the effect of region selection on the performance for each subject. On the top, their distributions are shown, along with their means (dashed lines) and medians (solid lines).

**Figure S5:**
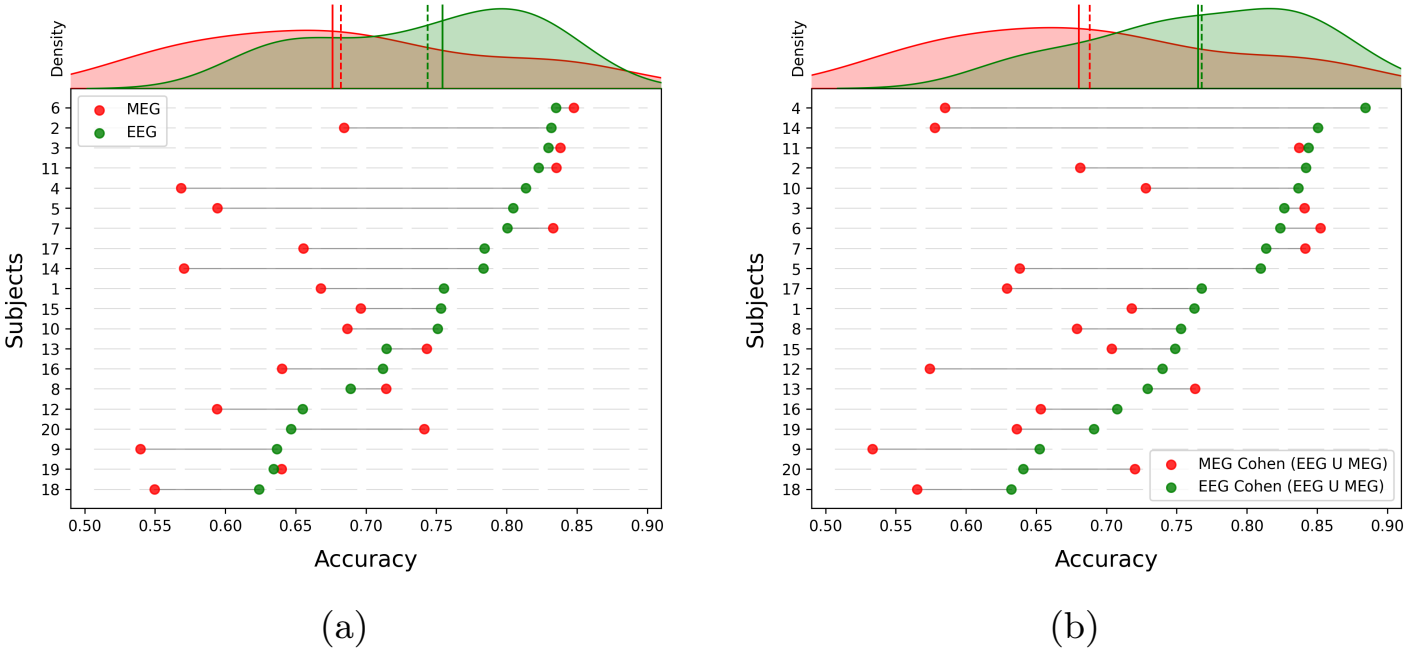
Lollipop plot comparing performances of single-modality pipelines. On the top, performance distributions are shown, along with their means (dashed lines) and medians (solid lines). (a) - ROI selection (based on Cohen’s d effect size) is performed independently for each modality. Subjects are ordered according to their EEG performance, obtained using the modality-specific ROI selection. (b) - ROI selection employed in this comparison is the Cohen (EEG ∪ MEG) criterion. Subjects are ordered according to their EEG performance, obtained using the shared Cohen (EEG ∪ MEG) ROI selection.

**Figure S6:**
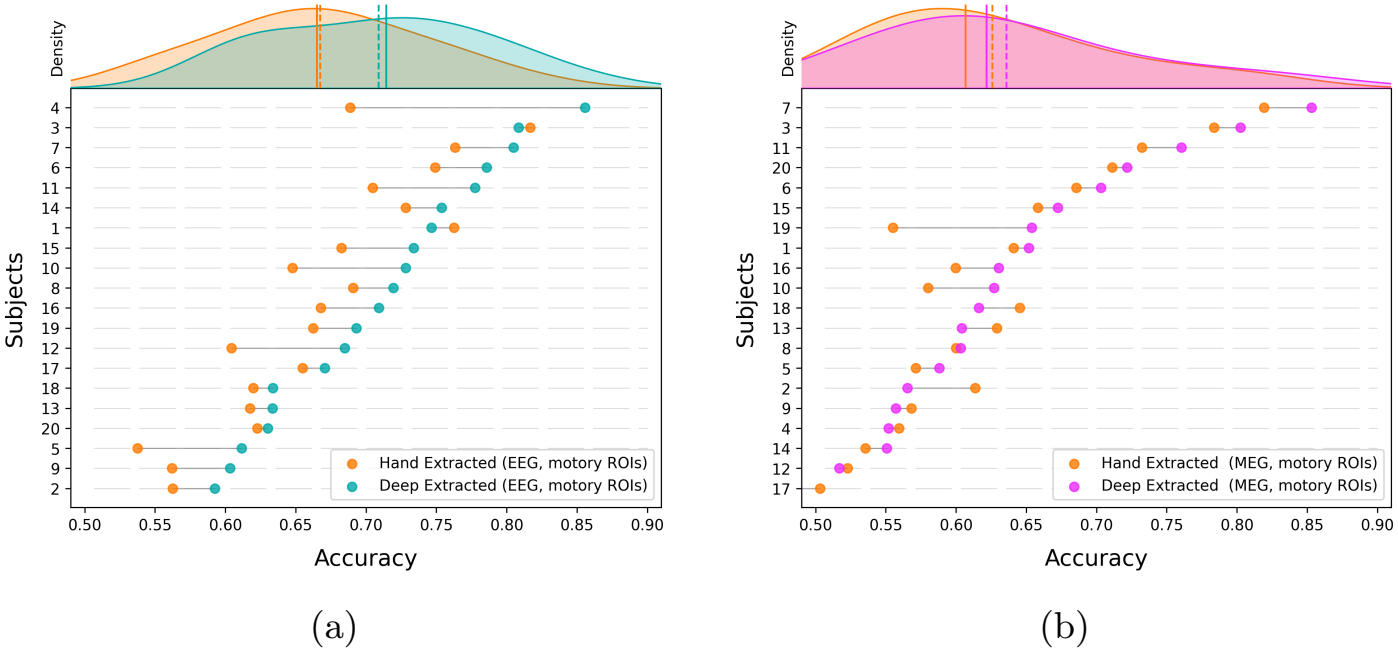
Lollipop plots comparing performance using encoder-extracted features versus PSD-based features, in the basic scenario of single modality and using sensorimotor ROIs. (a) - EEG modality case. Mean accuracy improves from 0.667 to 0.709 using our proposed method, while the median improves from 0.665 to 0.714. Here, subjects are ordered according to their EEG performance obtained with the encoder-decoder features. (b) - MEG modality case. Mean accuracy improves from 0.626 to 0.636 using our proposed method, while median improves from 0.607 to 0.622. Here, subjects are ordered according to their MEG performance obtained with the encoder-decoder features.

## Notes

### Competing Interest Statement

The authors have declared no competing interest.

### Summary of Updates

Update section 2.2.4: added specifications about dat splitting

