## Supplementary material for "FUSE: FUsing EEG–MEG in a Shared Embedding via self-supervised learning for BCI"

### 6 Supplementary

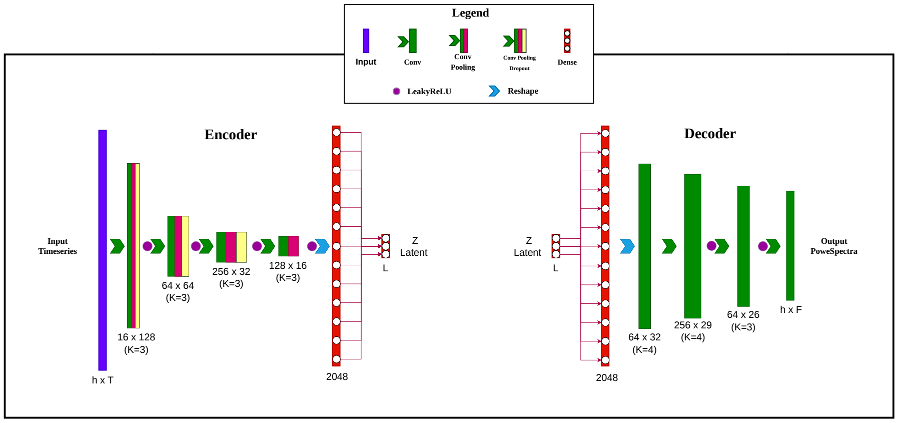


Figure S1: Architecture of the encoder-decoder network. Both the encoder and decoder employ a feedforward convolutional architecture. Each layer is represented as a rectangle, with its type indicated in the legend. The numbers below each layer indicate the corresponding output dimensions, while values in parentheses ($K = \cdot$) denote the kernel size of convolutional layers. All pooling operations are implemented using average pooling, and dropout layers use a dropout rate of 0.15. Padding is applied in the encoder convolutional layers to preserve power-of-two dimensions. No padding is used in the decoder. The encoder (left) consists of four convolutional layers followed by a fully connected layer, whose output defines the latent representation, the feature extracted. The decoder (right) receives this latent representation and estimates the spectral representation of the input time series through one fully connected layer followed by three convolutional layers.

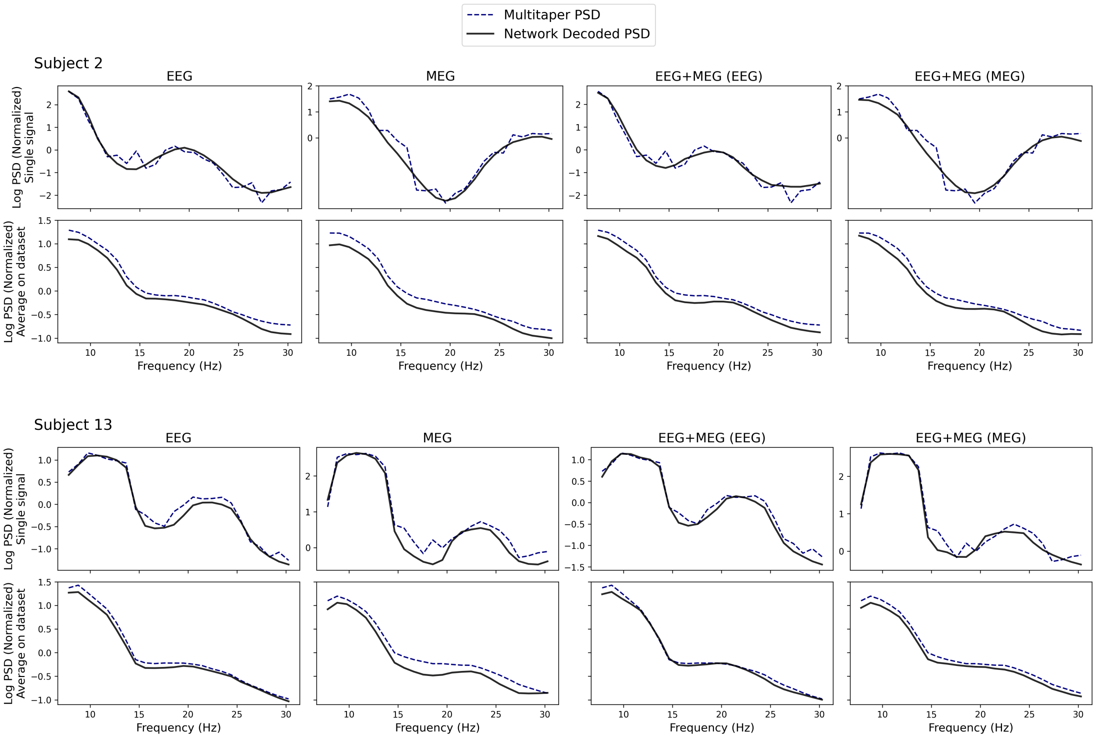


Figure S2: Ability of the encoder–decoder networks to reconstruct the target PSD. For conciseness, only Subjects 2 and 13 are shown, occupying the upper and lower halves of the figure, respectively. The first two columns show the reconstructions obtained from the single-modality networks, while the last two columns show the EEG and MEG reconstructions, respectively, obtained from the (same) two-modality network. For each subject, the first row shows an example of an individual PSD reconstruction, while the second row shows the average PSD across the *Val1* set, comparing the multitaper estimates with the corresponding network reconstructions.

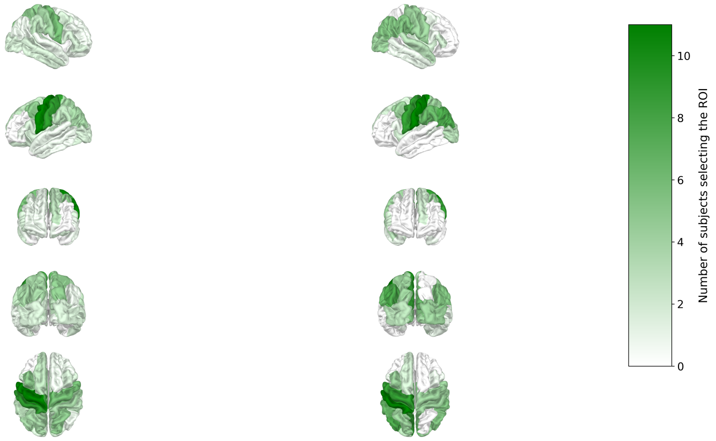


(a) EEG selection. (b) MEG selection.

Figure S3: ROIs selected by our criterion based on Cohen's d effect size, making use of all the data for each subject. The selection is performed independently for each subject. Each brain ROI is colored based on the number of subjects for whom that ROI is selected. The selection is performed both for EEG and MEG modalities, independently.

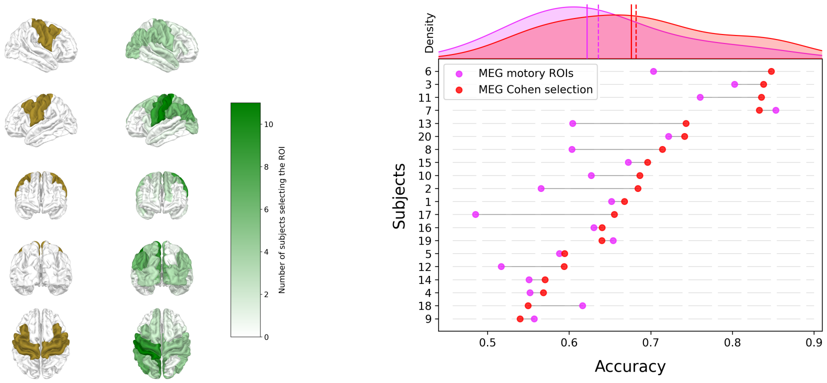


(a) (b)

Figure S4: Comparison of ROIs selected based on the Cohen-based selection ($n_{\text{roi}} = 8$) with sensorimotor ROIs and performance effects in the case of the MEG modality. Mean accuracy is 0.636 and 0.682, and median accuracy is 0.622 and 0.676, using sensorimotor ROIs and Cohen-selected ROIs, respectively. (a) - Brain plot comparing, in different visualizations, sensorimotor ROIs (in ochre) and ROIs selected by our criterion (in green) within the cross-validation loop for MEG. (b) - Lollipop plot showing the effect of region selection on the performance for each subject. On the top, their distributions are shown, along with their means (dashed lines) and medians (solid lines).

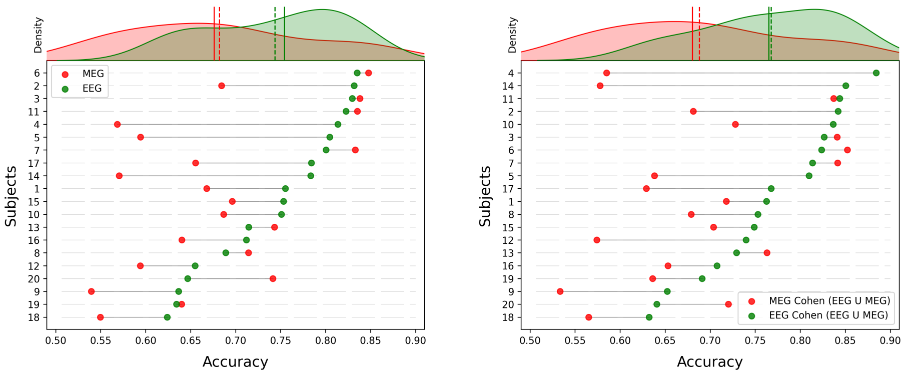


(a) (b)

Figure S5: Lollipop plot comparing performances of single-modality pipelines. On the top, performance distributions are shown, along with their means (dashed lines) and medians (solid lines). (a) - ROI selection (based on Cohen's d effect size) is performed independently for each modality. Subjects are ordered according to their EEG performance, obtained using the modality-specific ROI selection. (b) - ROI selection employed in this comparison is the Cohen (EEG $\cup$ MEG) criterion. Subjects are ordered according to their EEG performance, obtained using the shared Cohen (EEG $\cup$ MEG) ROI selection.

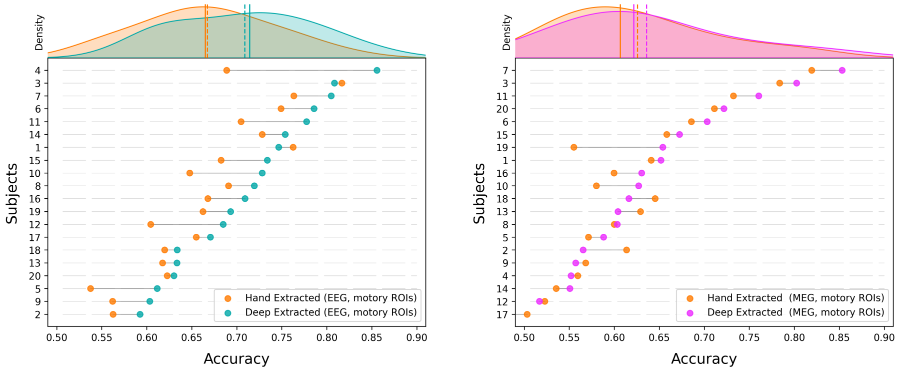


(a) (b)

Figure S6: Lollipop plots comparing performance using encoder-extracted features versus PSD-based features, in the basic scenario of single modality and using sensorimotor ROIs. (a) - EEG modality case. Mean accuracy improves from 0.667 to 0.709 using our proposed method, while the median improves from 0.665 to 0.714. Here, subjects are ordered according to their EEG performance obtained with the encoder-decoder features. (b) - MEG modality case. Mean accuracy improves from 0.626 to 0.636 using our proposed method, while median improves from 0.607 to 0.622. Here, subjects are ordered according to their MEG performance obtained with the encoder-decoder features.
